# Zika viruses of both African and Asian lineages cause fetal harm in a vertical transmission model

**DOI:** 10.1101/387118

**Authors:** Anna S. Jaeger, Reyes A. Murreita, Lea R. Goren, Chelsea M. Crooks, Ryan V. Moriarty, Andrea M. Weiler, Sierra Rybarczyk, Matthew R. Semler, Christopher Huffman, Andres Mejia, Heather A. Simmons, Michael Fritsch, Jorge E. Osorio, Shelby L. O’Connor, Gregory D. Ebel, Thomas C. Friedrich, Matthew T. Aliota

## Abstract

Congenital Zika virus (ZIKV) infection was first linked to birth defects during the American outbreak ^1–3^. It has been proposed that mutations unique to the Asian/American-genotype explain, at least in part, the ability of Asian/American ZIKV to cause congenital Zika syndrome (CZS) ^4,5^. Recent studies identified mutations in ZIKV infecting humans that arose coincident with the outbreak in French Polynesia and were stably maintained during subsequent spread to the Americas ^5^. Here we show that African ZIKV can infect and harm fetuses and that the S139N mutation that has been associated with the American outbreak is not essential for fetal harm. Our findings, in a vertical transmission mouse model, suggest that ZIKV will remain a threat to pregnant women for the foreseeable future, including in Africa, southeast Asia, and the Americas. Additional research is needed to better understand the risks associated with ZIKV infection during pregnancy, both in areas where the virus is newly endemic and where it has been circulating for decades.

## Main

Zika virus causes adverse pregnancy outcomes including fetal loss, developmental abnormalities, and neurological damage -- impacts that are collectively termed congenital Zika syndrome (CZS) ^6–9^. Why does CZS seem like a new complication when ZIKV has been circulating in Africa and Asia for decades? A provocative explanation for the recent appearance of CZS is that, during their geographic spread from Asia to the Americas, contemporary ZIKV strains acquired mutations that enhance neurovirulence. In several arboviruses, simple point mutations are known to result in changes in host range and/or the efficiency of infection and replication in key amplification hosts or vectors (see ^10^ for review). For example, a single serine-to-asparagine substitution in the transmembrane protein of ZIKV (prM; S139N) that is unique to the Asian/American lineage viruses is postulated to increase neurovirulence and contribute significantly to the microcephaly phenotype ^5^.

Yuan et al. ^5^ recently demonstrated that S139N substantially increased ZIKV infectivity in both human and mouse neural progenitor cells (NPCs), leading to restricted brain growth in an *ex-vivo* embryonic mouse brain model, as well as higher mortality rates in neonatal mice following intracranial (i.c.) inoculation. However, accumulating data suggest that in endemic areas, the virus has always been teratogenic ^11–14^. The degree to which the capacity to cause fetal harm is an emergent property unique to ZIKV circulating in the Americas remains an open question.

To more fully characterize the range of pathogenic outcomes of congenital ZIKV infection and to assess the role of S139N on an alternate genetic background, we engineered the reverse amino acid mutation (asparagine reverted to serine at residue 139 in the viral polyprotein) into the Puerto Rican ZIKV isolate PRVABC59 (ZIKV-PR-N139S). Prior to use in mice, we assessed viral infectivity and replication of ZIKV-PR-N139S *in vitro* using Vero cells. ZIKV-PR-N139S and a control virus derived from an infectious clone bearing the wild type ZIKV-PRVABC59 consensus sequence (ZIKV-PR-IC) gave similar growth curves (Fig. 1a). These results suggest that the “reverse substitution” N139S did not have a significant effect on either infectivity or replicative capacity *in vitro*. Next, to assess whether N139S, in the context of the PRVABC59 genome, decreased mortality in the neonatal mouse model, we inoculated one-day-old BALB/c mice i.c. with 10 PFU of either ZIKV-PR-IC; ZIKV-PR-N139S; a ZIKV strain isolated in Cambodia in 2010 (ZIKV-CAM; FSS 13025); a low-passage African ZIKV strain isolated in Senegal in 1984 (ZIKV-DAK; DAK AR 41524); or, as a control, phosphate-buffered saline (PBS). Surprisingly, and in contrast to results described by Yuan et al., i.c. inoculation of both ZIKV-DAK and ZIKV-CAM resulted in 100% mortality, whereas 80% of mice succumbed to ZIKV-PR-IC and 56% to ZIKV-PR-N139S by 28 days post inoculation (dpi; Fig. 1b). All strains caused significant mortality (log-rank test) as compared to the PBS-inoculated controls (ZIKV-CAM, ZIKV-DAK: *p*-value <0.0001; ZIKV-PR-IC: *p*-value = 0.002; ZIKV-PR-N139S: *p*-value = 0.016). ZIKV-PR-IC and ZIKV-PR-N139S mortality rates did not significantly differ (Fisher’s exact test *p*-value = 0.21). These results are consistent with other studies in pregnant animal models that have provided evidence of neurovirulence and fetal demise caused by ZIKV strains isolated before the American outbreak ^15,16^.

**Figure 1.**
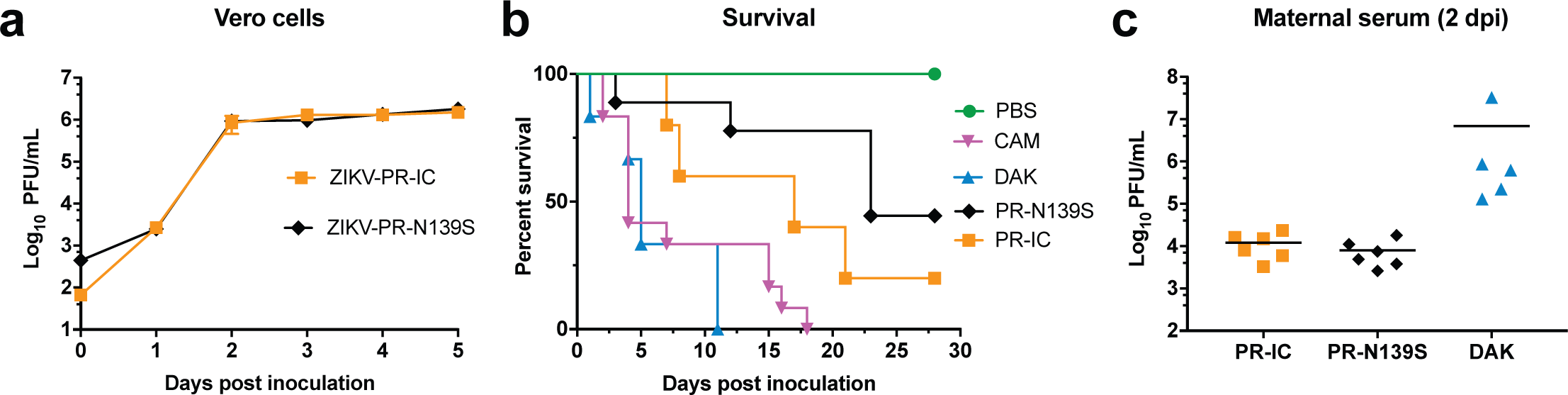
In vitro and in vivo characterization of ZIKV strains. **(a)**In vitro growth kinetics of ZIKV-PR-IC and mutant ZIKV-PR-N139S on Vero cells. Data points represent means of three replicates at each time point ± standard deviation. Cells were inoculated at an MOI of 0.01 PFU/cell. Titer was measured (PFU/ml) by plaque assay. Growth curves were not significantly different. **(b)**Neurovirulence phenotypes of different strains tested in neonatal mice. Neonatal BALB/c mice were intracranially inoculated with 10 PFU of ZIKV and mortality was recorded daily for 28 days. PBS: n=8; -CAM: n=12; -PR-IC: n=5; -PR-N139S: n=9; -DAK: n=6. All strains caused significant mortality when compared to PBS (log-rank test). Mortality was not significantly different between ZIKV-PR-IC and ZIKV-PR-N139S. *P <0.01; **P<0.001; ***P <0.0001; NS, not significant. **(c)**Time-mated Ifnar1-/-dams were inoculated with 103 PFU of ZIKV on E7.5 and maternal infection was confirmed by plaque assay on day 2 post inoculation. Titer did not significantly differ between strains (unpaired Student’s t-test).

The previous experiments establish the ability of each ZIKV strain tested to cause lethal infections in neonates, but direct intracranial inoculation does not fully recapitulate the events of a natural congenital infection. To better compare the abilities of these ZIKV strains to induce birth defects following vertical transmission, we used a previously established murine pregnancy model for ZIKV ^16,17^, in which dams lacking type I interferon signaling (*Ifnar1*^-/-^) were crossed with wildtype sires to produce heterozygous offspring. Because they have one intact *Ifnar1* haplotype, these offspring more closely resemble the immune status of human fetuses. Time-mated dams were inoculated subcutaneously in the footpad with 10^3^ PFU of ZIKV-PR-IC, ZIKV-PR-N139S, or ZIKV-DAK on embryonic day 7.5 (E7.5), corresponding to the mid-to-late first trimester in humans ^18^. We omitted ZIKV-CAM in the vertical transmission experiment due to previous experiments demonstrating its ability to cause fetal harm in this model ^16^. We collected serum samples from dams at peak viremia (2 dpi) to confirm infection and to sequence viral populations replicating *in vivo*. All dams were productively infected without significant differences in titer (Student’s t-test) between treatment groups (PR-IC vs. -PR-N139S: *p*-value = 0.33, *t*-value = 1.03, df = 10; -PR-IC vs. -DAK: *p*-value = 0.27, *t*-value = 1.19, df = 9; -PR-N139S vs. -DAK: *p*-value = 0.27, *t*-value = 1.19, df = 9; Fig. 1c). Deep sequencing of virus populations replicating in maternal serum confirmed that the N139S mutation was stably maintained *in vivo* (Table 1). Dams were monitored daily for clinical signs until time of necropsy. Overt clinical signs were only evident in ZIKV-DAK-inoculated dams and included hunched posture, ruffled fur, and hind limb paralysis indicative of neurotropism. All ZIKV-DAK infected dams met euthanasia criteria at time of necropsy on E14.5.

**Table 1.** Deep sequencing of virus populations replicating in maternal serum. Serum samples from pregnant mice infected with ZIKV-PR-IC, ZIKV-PR-N139S, and ZIKV-DAK were deep sequenced in duplicate (R1, R2) and analyzed with the “Zequencer 2017” workflow to confirm the expected residue at position 139. The major codon found at residue 139 for each sample is shown in red.

|  |  | Major codon<br>frequency at<br>residue 139 (%) | Subsampled coverage<br>depth at residue 139<br>(# of reads) |
| --- | --- | --- | --- |
| **ZIKV-PRVABC59 (KU501215)** | 130<br>S A Y Y M Y L D R **N** D<br>140 | **R1/R2** | **R1/R2** |
| ZIKV-PR-IC-1 | . . . . . . . . . . . | 98.9/98.7 | 2000/149 |
| ZIKV-PR-IC-2 | . . . . . . . . . . . | 99.2/94.5 | 2000/110 |
| ZIKV-PR-IC-3 | . . . . . . . . . . . | 97.1/100.0 | 1999/16 |
| ZIKV-PR-IC-4 | . . . . . . . . . . . | 99.8/99.3 | 2000/1023 |
| ZIKV-PR-IC-5 | . . . . . . . . . . . | 99.8/99.9 | 2000/2000 |
| ZIKV-PR-IC-6 | . . . . . . . . . . . | 99.9/99.7 | 2000/1715 |
| ZIKV-PR-N139S-1 | . . . . . . . . . **S** . | 99.8/99.7 | 2000/1502 |
| ZIKV-PR-N139S-2 | . . . . . . . . . **S** . | 99.9/99.7 | 2000/902 |
| ZIKV-PR-N139S-3 | . . . . . . . . . **S** . | 100.0/99.8 | 1440/2000 |
| ZIKV-PR-N139S-4 | . . . . . . . . . **S** . | 99.7/99.7 | 2000/2000 |
| ZIKV-PR-N139S-5 | . . . . . . . . . **S** . | 99.6/98.9 | 2000/709 |
| ZIKV-PR-N139S-6 | . . . . . . . . . **S** . | 99.6/99.7 | 2000/1585 |
| **ZIKV-DAK AR 41524 (KX601166.2)** | 130<br>S A Y Y M Y L D R **S** D<br>140 | **R1/R2** | **R1/R2** |
| ZIKV-DAK-1 | . . . . . . . . . . . | 100.0/100.0 | 2000/2000 |
| ZIKV-DAK-2 | . . . . . . . . . . . | 100.0/100.0 | 2000/2000 |
| ZIKV-DAK-3 | . . . . . . . . . . . | 100.0/99.9 | 2000/2000 |
| ZIKV-DAK-4 | . . . . . . . . . . . | 99.9/99.9 | 2000/2000 |
| ZIKV-DAK-5 | . . . . . . . . . . . | 99.9/100.0 | 2000/2000 |

Next, to assess fetal outcomes, ZIKV-inoculated dams were sacrificed at E14.5 and gross examination of each conceptus (both fetus and placenta, when possible) revealed overt differences among fetuses within pregnancies and with uninfected counterparts. In general, fetuses appeared either grossly normal or abnormal, defined as being in one of the four stages of embryo resorption (Fig. 2a, c)^19^. At time of necropsy, we observed high rates of resorption in both ZIKV-PR-IC- and ZIKV-PR-N139S-infected pregnancies. The proportion of abnormal fetuses for the two strains did not differ significantly (53.2%, 67.3%, Fisher’s exact test *p*-value = 0.21). In contrast, ZIKV-DAK-infected pregnancies resulted in 100% resorption of fetuses (Fig 2a). Only fetuses that appeared grossly normal were included for measurement of crown-rump length (CRL) to provide evidence for intrauterine growth restriction (IUGR). There was a modest reduction in size in grossly normal ZIKV-PR-IC fetuses. Mean CRL did not differ significantly (Student’s t-test) between fetuses of ZIKV-PR-IC- or PBS-inoculated dams (*p*-value = 0.22, *t-*value = 1.23, df = 54), whereas there was a statistically significant (Student’s t-test) reduction in mean CRL between fetuses whose dams were inoculated with ZIKV-PR-IC vs. ZIKV-PR-N139S (*p*-value <0.0001, *t*-value = 5.42, df = 34; Fig 2b). This lack of evidence of severe IUGR for ZIKV-PR-IC is contrary to other studies in which fetuses developed severe IUGR ^16,17,20^; however, this may be due to differences in timing of challenge and necropsy ^21^, virus strain, or a different standard for characterizing grossly normal fetuses compared to those in a stage of resorption at a later embryonic age.

**Figure 2:**
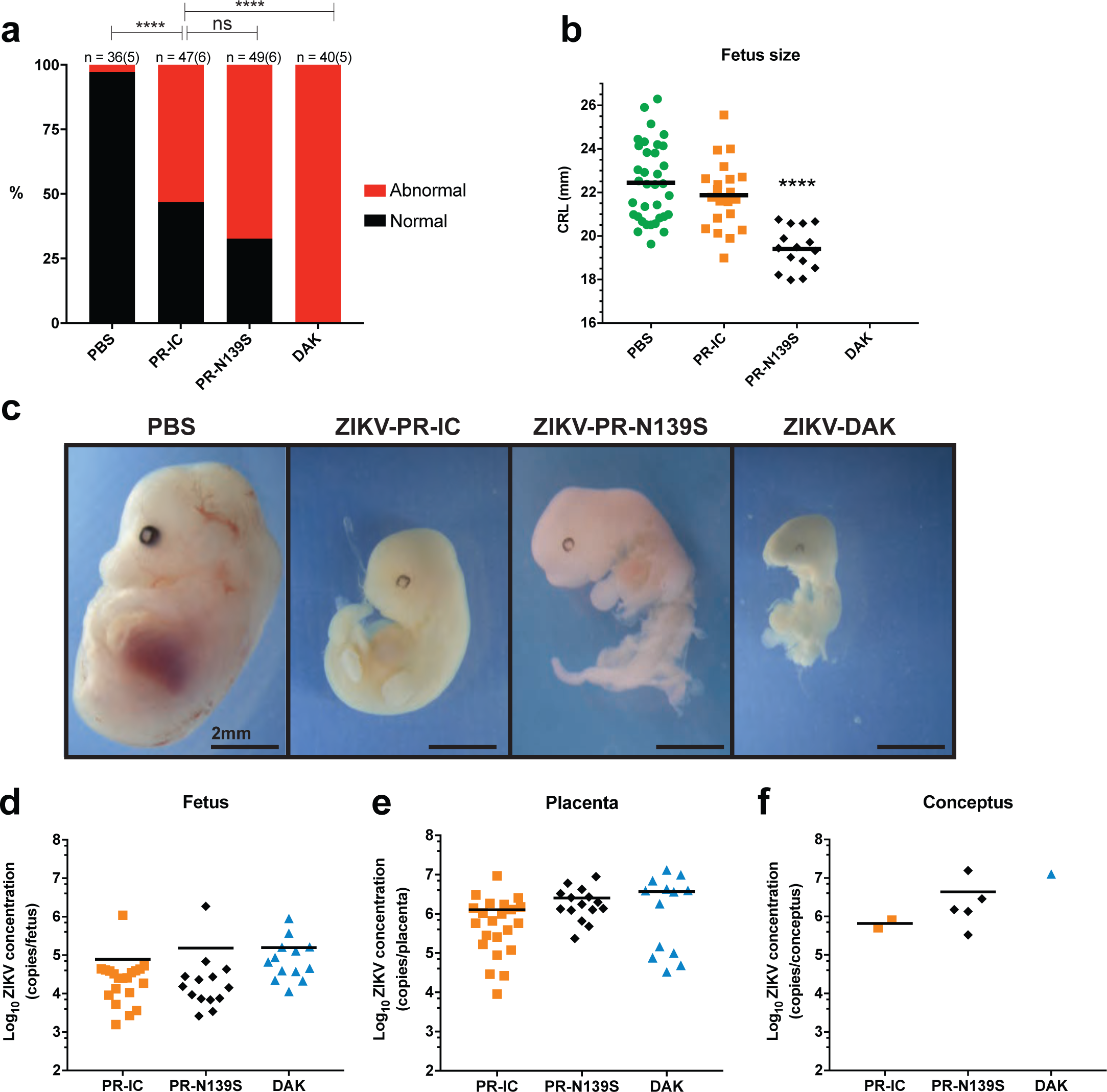
Fetal outcomes after maternal infection with ZIKV strains. **(a)**Rate of grossly normal (black) versus abnormal (red) fetuses at E14.5 after maternal infection at E7.5. An abnormal fetus was defined as in any of the four stages of resorption. Data presented are for individual fetuses from 5-6 litters per treatment group. The n for each group is indicated above each bar. ****P<0.0001; NS, not significant (Fisher’s exact test). **(b)**Fetus size as assessed by crown-rump length (CRL) in mm using ImageJ software. CRL was only measured for fetuses determined to be grossly normal at E14.5. CRL of ZIKV-PR-IC fetuses did not differ significantly from PBS. There were no grossly normal fetuses for measurement from ZIKV-DAK litters. ***P<0.0001; NS, not significant (unpaired Student’s t-test). **(c)**Representative images of fetuses on E14.5 from each treatment group. Scale bar, 2 mm. PBS fetus characterized as normal. -PR-IC, -PR-N139S, -DAK fetuses characterized as abnormal. **(d-f)**Viral burdens were measured by qRT-PCR assay from individual homogenized placentas **(d)**, fetuses **(e)**, and concepti (when fetus and placenta could not be separated due to severe resorption) **(f)**. Symbols represent individual placenta, fetus, or conceptus from 3-5 independent experiments for each treatment group. Viral burdens were not significantly different between treatment groups (unpaired Student’s t-test).

To confirm vertical transmission of ZIKV to the developing conceptus, viral loads were measured from representative placentas and fetuses from each litter of all treatment groups by quantitative RT-PCR (Fig 2d-f). vRNA was detected in all fetuses and placentas that were tested. Viral loads were significantly higher (Student’s t-test) in placentas than in fetuses (*p*-value <0.0001, *t*-value =5.04, df = 95; Fig 2d-e), whereas viral loads were not significantly different between groups infected with different viruses nor among littermates within the same litter (-PR-IC vs. -PR-N139S: *p*-value = 0.61, *t*-value = 0.52, df = 17.3; -PR-IC vs. -DAK: *p*-value = 0.36, *t*-value = 0.93, df = 25.5; -PR-N139S vs. -DAK: *p*-value = 0.97, *t*-value = 0.04, df = 19.2; Fig. 1c). Detection of ZIKV RNA in grossly normal fetuses does not preclude the possibility that pathology may develop later in pregnancy or even postnatally, consistent with reports from humans that the effects of *in utero* exposure may not be evident at birth ^22^.

To better understand the impact of *in utero* ZIKV infection, tissues of the developing placenta and decidua were evaluated microscopically. In PBS-inoculated dams, we observed normal decidua, junctional zone, and labyrinth with normal maternal and fetal blood spaces (Fig 3a-c). In contrast, ZIKV-inoculated dams displayed varying degrees of placental pathology, including vascular injury involving maternal and/or fetal vascular spaces, infarction (obstructed blood flow), necrosis, inflammation, and hemorrhage (Fig 3d-f). There also were clear strain-specific differences in the amount of placental pathology, with ZIKV-DAK displaying the most severe histologic phenotype, consistent with gross observations (Fig 3g-i).

**Figure 3:**
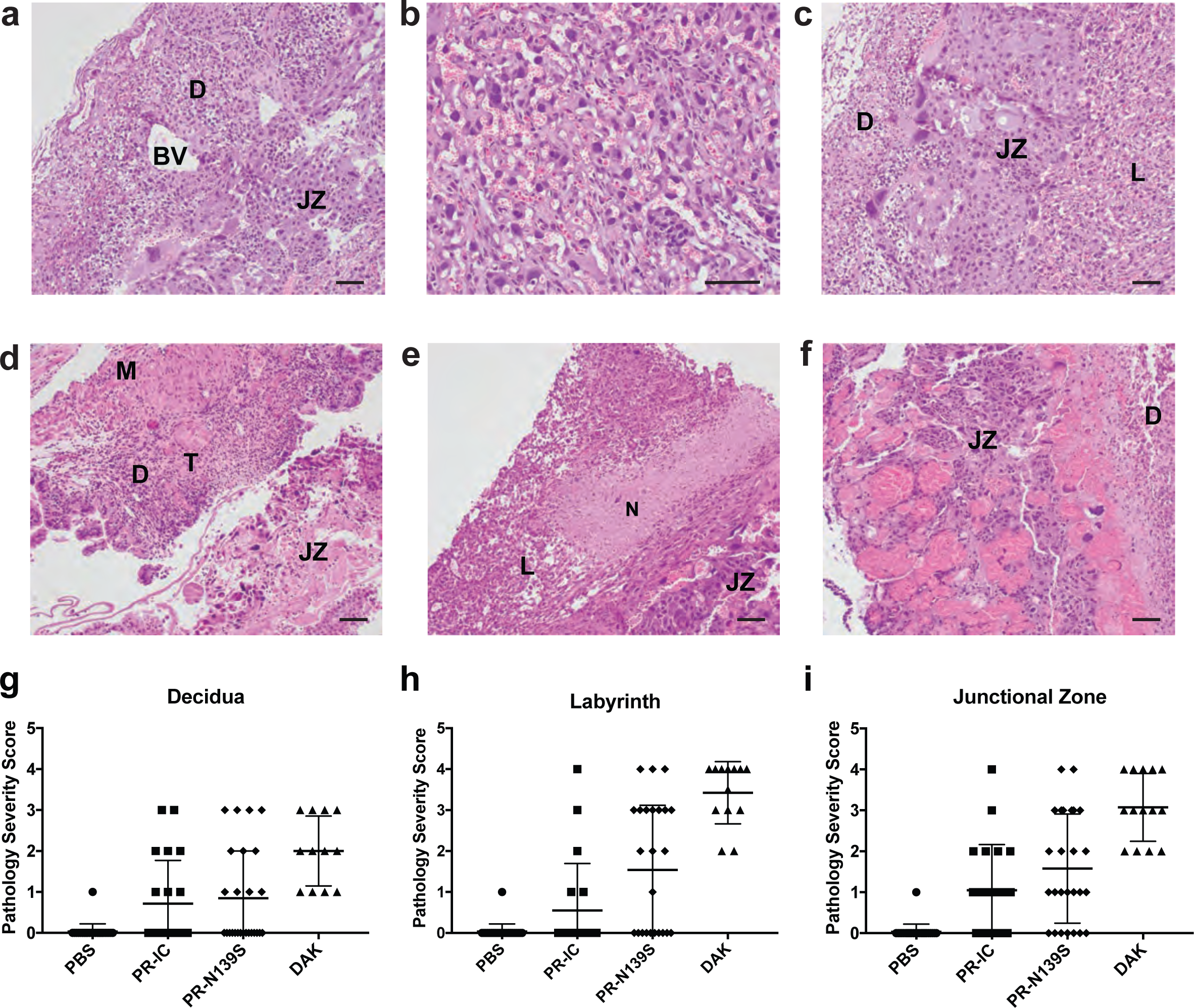
Placenta histopathology analysis: Hematoxylin and eosin (H&E) staining of placenta and fetus. **(a-c)**Normal histologic features of each placental zone (decidual layer (D), labyrinth layer (L), and junctional zone (JZ)) from concepti from dams inoculated with PBS. The decidua is the maternally-derived endometrial lining of the uterus, and the fetus-derived placenta is composed of the JZ and L, where nutrient exchange occurs between the maternal and fetal blood. In L fetal blood spaces contain nucleated red blood cells and maternal spaces contain only red cells without nuclei. BV, normal decidual blood vessels. **(d-f)**Severe histopathologic injury patterns for each zone from placenta from ZIKV-inoculated dams. **(d)**Myometrium (M) and decidua (D) from a ZIKV-PR-IC placenta with increased inflammation, multiple thrombi (T) in the decidua, and a necrotic JZ. **(e)**L from a ZIKV-DAK placenta with focal necrosis (N), lack of blood in most vascular spaces, and numerous degenerating cells. **(f)**JZ from a ZIKV-DAK placenta with markedly dilated blood vessels, focal thrombi, and a layer of necrosis at the interface with the decidua. **(g-i)**The degree of placental pathology was rated on a scale of 0-4: zero represents normal histologic features and 4 represents the most severe features observed. Each zone of the placenta was scored individually for general overall pathology, amount of inflammation, and amount of vascular injury with a consensus score for each placenta derived from three independent pathologists. Only ‘General’ scores are shown because they were representative of the ‘inflammation’ and ‘vascular injury’ categories and do not differ significantly from ‘general’. Error bars represent standard deviation of the mean. Data are representative of 3-5 independent experiments for each treatment group. Scale bar, 50 µm.

Together our data show that infection with ZIKV isolates of either the African or Asian lineages during pregnancy can lead to fetal harm, with varying levels of damage to maternal, placental, and fetal tissues, frequently including death of the developing fetus. Likewise, intracranial inoculation of neonatal mice confirmed a similar neurovirulence phenotype across ZIKV lineages. The observation that a low-passage African ZIKV isolate can cause severe fetal harm suggests that, for decades, ZIKV could have been causing pregnancy losses and birth defects, which were either undiagnosed or attributed to other causes. If this hypothesis is correct, CZS is not a new syndrome caused by a recently emerged ZIKV variant, but rather an old entity that was only recognized in the large-scale American ZIKV outbreak that began in 2014-15. These results provide compelling motivation to re-evaluate hypotheses explaining the emergence of CZS. A lack of thorough surveillance, paired with myriad co-circulating febrile illnesses, make understanding both the past and current prevalence of gestational ZIKV infection and any resulting fetal outcomes in Africa challenging ^23–25^. Recent seroprevalence studies have now identified low, but consistent circulation of ZIKV in several African countries ^26–29^, undermining the hypothesis that herd immunity protects against CZS and indicating a large population potentially at risk. Accurate assessment of the risk posed by ZIKV infection to pregnant women and their babies in both Africa and southeast Asia should be a priority.

## Methods

### Ethical approval

This study was approved by the University of Wisconsin-Madison Institutional Animal Care and Use Committee (Animal Care and Use Protocol Number V5519).

### Cells and Viruses

African Green Monkey kidney cells (Vero; ATCC #CCL-81) were maintained in Dulbecco’s modified Eagle medium (DMEM) supplemented with 10% fetal bovine serum (FBS; Hyclone, Logan, UT), 2 mM L-glutamine, 1.5 g/L sodium bicarbonate, 100 U/ml penicillin, 100 µg/ml of streptomycin, and incubated at 37°C in 5% CO_2_. *Aedes albopictus* mosquito cells (C6/36; ATCC #CRL-1660) were maintained in DMEM supplemented with 10% fetal bovine serum (FBS; Hyclone, Logan, UT), 2 mM L-glutamine, 1.5 g/L sodium bicarbonate, 100 U/ml penicillin, 100 µg/ml of streptomycin, and incubated at 28°C in 5% CO_2_. The cell lines were obtained from the American Type Culture Collection, were not further authenticated, and were not specifically tested for mycoplasma. ZIKV strain PRVABC59 (ZIKV-PR; GenBank:KU501215), originally isolated from a traveler to Puerto Rico in 2015 with three rounds of amplification on Vero cells, was obtained from Brandy Russell (CDC, Ft. Collins, CO). ZIKV-PR served as the backbone for the reverse genetic platform developed by Weger-Lucarelli et al. ^30^ upon which the single-amino acid substitution-N139S-was introduced. ZIKV strain DAK AR 41524 (ZIKV-DAK; GenBank:KX601166) was originally isolated from *Aedes luteocephalus* mosquitoes in Senegal in 1984, with a round of amplification on *Aedes pseudocutellaris* cells, followed by amplification on C6/36 cells, followed by two rounds of amplification on Vero cells, was obtained from BEI Resources (Manassas, VA). Virus stocks were prepared by inoculation onto a confluent monolayer of C6/36 mosquito cells. ZIKV strain FSS 13025 (ZIKV-CAM; GenBank:JN860885), originally isolated from a child in Cambodia in 2010 with three rounds of amplification on Vero cells, was obtained by Brandy Russell (CDC, Ft. Collins, CO). Virus stocks were prepared by inoculation onto a confluent monolayer of Vero cells.

### Generation of ZIKV prM mutant

An infectious clone for ZIKV-PR was constructed as previously described ^30^. Infectious-clone derived virus (ZIKV-PR-IC) was recovered following electroporation of *in vitro* transcribed RNA into Vero cells. To engineer the N139S mutation into the ZIKV genome, the corresponding single-amino acid substitution was introduced into the ZIKV-PR-IC using the IVA cloning method ^31^. The infectious clone plasmids were linearized by restriction endonuclease digestion, PCR purified, and ligated with T4 DNA ligase. From the assembled fragments, capped T7 RNA transcripts were generated, and the resulting RNA was electroporated into Vero cells. Infectious virus was harvested when 50-75% cytopathic effects were observed (6 days post transfection; ZIKV-N139S). Viral supernatant then was clarified by centrifugation and supplemented to a final concentration of 20% fetal bovine serum and 10 mM HEPES prior to freezing and storage as single use aliquots. Titer was measured by plaque assay on Vero cells as described in a subsequent section. We deep sequenced all of our challenge stocks (both wildtype and infectious clone-derived viruses) to verify the expected origin and amino acid at residue 139 (see details in a section below). All ZIKV stocks had the expected amino acid at residue 139: ZIKV-PR-IC (N), ZIKV-DAK (S), ZIKV-CAM (S), ZIKV-PR-N139S (S). Importantly, no single nucleotide polymorphisms were detected at residue 139 at a frequency greater than 1%, nor did we detect evidence of Dezidougou virus, an insect-specific *Negevirus* present in some ZIKV DAK AR 41524 stocks.

### Plaque assay

All ZIKV screens from mouse tissue and titrations for virus quantification from virus stocks were completed by plaque assay on Vero cell cultures. Duplicate wells were infected with 0.1 ml aliquots from serial 10-fold dilutions in growth media and virus was adsorbed for one hour. Following incubation, the inoculum was removed, and monolayers were overlaid with 3 ml containing a 1:1 mixture of 1.2% oxoid agar and 2X DMEM (Gibco, Carlsbad, CA) with 10% (vol/vol) FBS and 2% (vol/vol) penicillin/streptomycin. Cells were incubated at 37 °C in 5% CO_2_ for four days for plaque development. Cell monolayers then were stained with 3 ml of overlay containing a 1:1 mixture of 1.2% oxoid agar and 2X DMEM with 2% (vol/vol) FBS, 2% (vol/vol) penicillin/streptomycin, and 0.33% neutral red (Gibco). Cells were incubated overnight at 37 °C and plaques were counted.

### Viral RNA isolation

Viral RNA was extracted from sera using the Viral Total Nucleic Acid Kit (Promega, Madison, WI) on a Maxwell 48 RSC instrument (Promega, Madison, WI). Viral RNA was isolated from homogenized tissues using the Maxwell 48 RSC Viral Total Nucleic Acid Purification Kit (Promega, Madison, WI) on a Maxwell 48 RSC instrument. Each tissue was homogenized using PBS supplemented with 20% FBS and penicillin/streptomycin and a tissue tearor variable speed homogenizer. Supernatant was clarified by centrifugation and the isolation was continued according to the Maxwell 48 RSC Viral Total Nucleic Acid Purification Kit protocol, and samples were eluted into 50 µl RNase free water. RNA was then quantified using quantitative RT-PCR. Viral load data from serum are expressed as vRNA copies/mL. Viral load data from tissues are expressed as vRNA copies/tissue.

### Quantitative reverse transcription PCR (QRT-PCR)

For ZIKV-PR, vRNA from serum and tissues was quantified by QRT-PCR using primers with a slight modification to those described by Lanciotti et al. to accommodate African lineage ZIKV sequences ^32^. The modified primer sequences are: forward 5’-CGYTGCCCAACACAAGG-3’, reverse 5’-CACYAAYGTTCTTTTGCABACAT-3’, and probe 5’-6fam-AGCCTACCTTGAYAAGCARTCAGACACYCAA-BHQ1-3’. The RT-PCR was performed using the SuperScript III Platinum One-Step Quantitative RT-PCR system (Invitrogen, Carlsbad, CA) on a LightCycler 480 instrument (Roche Diagnostics, Indianapolis, IN). The primers and probe were used at final concentrations of 600 nm and 100 nm respectively, along with 150 ng random primers (Promega, Madison, WI). Cycling conditions were as follows: 37°C for 15 min, 50°C for 30 min and 95°C for 2 min, followed by 50 cycles of 95°C for 15 sec and 60°C for 1 min. Viral RNA concentration was determined by interpolation onto an internal standard curve composed of seven 10-fold serial dilutions of a synthetic ZIKV RNA fragment based on a ZIKV strain derived from French Polynesia that shares >99% similarity at the nucleotide level to the Puerto Rican strain used in the infections described in this manuscript.

### *In vitro* viral replication

Six-well plates containing confluent monolayers of Vero cells were infected with virus (ZIKV-PR-IC or ZIKV-PR-N139S), in triplicate, at multiplicity of infection (MOI) of 0.01 PFU/ml. After one hour of adsorption at 37º, the inoculum was removed and the cultures were washed three times. Fresh media were added and Vero cell cultures were incubated for 5 days 37°C, with aliquots removed daily, diluted 1:10 in culture media, and stored at −80°C. Viral titers at each time point were determined by plaque titration on Vero cells and viral loads were determined by QRT-PCR.

### Mice

Female *Ifnar-/-* mice on the C57BL/6 background were bred in the pathogen-free animal facilities of the University of Wisconsin-Madison Mouse Breeding Core within the School of Medicine and Public Health. Male C57BL/6 mice were purchased from Jackson Laboratories. Untimed, pregnant BALB/c mice were purchased from Charles River.

### Subcutaneous inoculation

All pregnant dams were between six and eight weeks of age. Littermates were randomly assigned to infected and control groups. Matings between female *Ifnar1^-/-^* dams and wildtype sires were timed by checking for the presence of a vaginal plug, indicating a gestational age E0.5. At embryonic day E7.5, dams were inoculated in the left, hind foot pad with 10^3^ PFU of ZIKV in 25 µl of sterile PBS or with 25 µl of sterile PBS alone to serve as experimental controls. All animals were closely monitored by laboratory staff for adverse reactions and signs of disease. A single sub-mandibular blood draw was performed 2 days post inoculation and serum was collected to verify viremia. Mice were were humanely euthanized and necropsied at E14.5.

### Mouse Necropsy

Following inoculation with ZIKV or PBS, mice were sacrificed at E14.5. Tissues were carefully dissected using sterile instruments that were changed between each mouse to minimize possible cross contamination. For all mice, each organ/neonate was evaluated grossly *in situ*, removed with sterile instruments, placed in a sterile culture dish, photographed, and further processed to assess viral burden and tissue distribution or banked for future assays. Briefly, uterus was first removed, photographed, and then dissected to remove each individual conceptus (i.e, fetus and placenta when possible). Fetuses and placentas were either collected in PBS supplemented with 20% FBS and penicillin/streptomycin (for plaque assays) or fixed in 4% PFA for imaging. Crown-rump length was measured by tracing distance from crown of head to end of tail using ImageJ. Infection-induced resorbed fetuses (~61%) were excluded from measurement analyses because they were unlikely to survive if the pregnancy was allowed to progress to term.

### Histology

Tissues were fixed in 4% paraformaldehyde for 24 hours and transferred into cold, sterile DPBS until alcohol processed and embedded in paraffin. Paraffin sections (5 μm) were stained with hematoxylin and eosin (H&E). Pathologists were blinded to gross pathological findings when tissue sections were evaluated microscopically. The degree of pathology at the maternal-fetal interface was rated on a scale of 0-4: 0 – no lesions (normal); 1 – mild changes; 2 – mild to moderate changes; 3 – moderate to severe changes; 4 – severe. The final scores were determined as a consensus score of three independent pathologists. For each zone in the placenta (myometrium, decidua, junctional zone, labyrinth, and chorionic plate/membranes) a ‘General’ overall score was determined, a score for the amount of ‘Inflammation’, and a score for direct ‘Vascular Injury’. The ‘General’ score was based on an interpretation of the overall histopathologic findings in each placenta, which included features of necrosis, infarction, hemorrhage, mineralization, vascular injury, and inflammation. The ‘Inflammation’ score quantified the amount of inflammation in that layer. The ‘Vascular Injury’ score assessed vascular wall injury (fibrinoid necrosis, endothelial swelling), dilatation of the vessels or spaces, and intraluminal thrombi. The myometrial layer representing the uterine wall and the chorionic plate/membranes were often not present in histologic sections and therefore meaningful comparisons between strains could not be assessed. The decidual layer (maternal in origin), the junctional zone composed of fetal giant cells and spongiotrophoblast, and the labyrinth layer (the critical layer for gas and nutrient exchange between the fetal and maternal vascular systems) were scored. Photomicrographs were obtained using a bright light microscope Olympus BX43 and Olympus BX46 (Olympus Inc., Center Valley, PA) with attached Olympus DP72 digital camera (Olympus Inc.) and Spot Flex 152 64 Mp camera (Spot Imaging), and captured using commercially available image-analysis software (cellSens DimensionR, Olympus Inc. and spot software 5.2).

### Intracranial inoculation

To test ZIKV strain neurovirulence, one-day-old BALB/c mice were intracranially (ic) inoculated at the lambda point with 10 PFU of virus or PBS alone. Following ic inoculation, mice were monitored twice daily for 28 days. Average survival time and percent mortality were calculated.

### Deep Sequencing

Virus populations replicating in mouse sera were sequenced in duplicate using a method adapted from Quick et. al. ^33^. Viral RNA was isolated from mouse sera using the Maxwell 16 Total Viral Nucleic Acid Purification kit, according to manufacturer’s protocol. Viral RNA then was subjected to RT-PCR using the SuperScript IV Reverse Transcriptase enzyme (Invitrogen, Carlsbad, CA). Input viral RNA was 10^6^ viral RNA templates per cDNA reaction. For sera from mice infected with ZIKV-PR-IC and ZIKV-PR-N139S, the cDNA was then split into two multi-plex PCR reactions using the PCR primers described in Quick et. al with the Q5® High-Fidelity DNA Polymerase enzyme (New England Biolabs®, Inc., Ipswich, MA). For sera from mice infected with ZIKV-DAK, the cDNA was amplified in a PCR reaction for sequencing of a single amplicon with ZIKV-DAK specific primers (forward 5’-ACCTTGCTGCCATGTTGAGA-3’, reverse 5’ CCGTACACAACCCAAGTCGA-3’) using Q5® High-Fidelity DNA Polymerase (New England Biolabs®, Inc., Ipswich, MA). PCR products were tagged with the Illumina TruSeq Nano HT kit and sequenced with a 2 x 250 kit on an Illumina MiSeq.

A vial of the viral stocks used for primary challenge (ZIKV-PR-IC, ZIKV-PR-N139S, ZIKV-DAK, ZIKV-CAM), were each deep sequenced by preparing libraries of fragmented double-stranded cDNA using methods similar to those previously described ^34^. Briefly, the sample was centrifuged at 5000 rcf for five minutes. The supernatant was then filtered through a 0.45-μm filter. Viral RNA was isolated using the QIAamp MinElute Virus Spin Kit (Qiagen, Germantown, MD), omitting carrier RNA. Eluted vRNA was then treated with DNAse I. Double-stranded DNA was prepared with the Superscript Double-Stranded cDNA Synthesis kit (Invitrogen, Carlsbad, CA) and priming with random hexamers. Agencourt Ampure XP beads (Beckman Coulter, Indianapolis, IN) were used to purify double-stranded DNA. The purified DNA was fragmented with the Nextera XT kit (Illumina, Madison, WI), tagged with Illumina-compatible primers, and then purified with Agencourt Ampure XP beads. Purified libraries were then sequenced with 2 x 300 bp kits on an Illumina MiSeq.

### Sequence analysis

Amplicon data were analyzed using a workflow we term “Zequencer 2017” (https://bitbucket.org/dhoconno/zequencer/src). Briefly, R1 and R2 fastq files from the paired-read Illumina miSeq dataset were merged, trimmed, and normalized using the bbtools package (http://jgi.doe.gov/data-and-tools/bbtools) and Seqtk (https://github.com/lh3/seqtk). Bbmerge.sh was used to merge reads, and to trim primer sequences by setting the forcetrimleft parameter 22. All other parameters are set to default values. These reads were then mapped to the reference amplicon sequences with BBmap.sh. Reads substantially shorter than the amplicon were filtered out by reformat.sh (the minlength parameter was set to the length of the amplicon minus 60). Seqtk was used to subsample to 1000 reads per amplicon. Quality trimming was performed on the fastq file of normalized reads by bbmap’s reformat.sh (qtrim parameter set to ‘lr’, all other parameters set to default). Novoalign (http://www.novocraft.com/products/novoalign/) was used to map each read to the appropriate ZIKV reference sequence: ZIKV-PRVABC59 KU501215, ZIKV DAK AR 41524 KX601166, ZIKV FSS13025 JN860885. Novoalign’s soft clipping feature was turned off by specifying the parameter “-o FullNW”. Approximate fragment length was set to 300bp, with a standard deviation of 50. We used Samtools to map, sort, and create an mpileup of our reads (http://samtools.sourceforge.net/). Samtools’ base alignment quality (BAQ) computation was turned off; otherwise, default settings were used. SNP calling was performed with VarScan’s mpileupcns function (http://varscan.sourceforge.net/). The minimum average quality was set to 30; otherwise, default settings were used. VCF files were annotated using SnpEff ^35^. Accurate calling of end-of-read SNPs are a known weakness of current alignment algorithms ^36^; in particular, Samtools’ BAQ computation feature is known to be a source of error when using VarScan (http://varscan.sourceforge.net/germline-calling.html). Therefore, both Novoalign’s soft clipping feature and Samtools’ BAQ were turned off to increase the accuracy of SNP calling for SNPs occurring at the end of a read.

Viral stock sequences were analyzed using a modified version of the viral-ngs workflow developed by the Broad Institute (http://viral-ngs.readthedocs.io/en/latest/description.html) implemented in DNANexus and using bbmap local alignment in Geneious Pro (Biomatters, Ltd., Auckland, New Zealand). Briefly, using the viral-ngs workflow, host-derived reads that map to a human sequence database and putative PCR duplicates are removed. The remaining reads were loaded into Geneious Pro and mapped to NCBI Genbank Zika virus reference sequences using bbmap local alignment. Mapped reads were aligned using Geneious global alignment and the consensus sequence was used for intrasample variant calling. Variants were called that fit the following conditions: have a minimum p-value of 10e-60, a minimum strand bias of 10e-5 when exceeding 65% bias, and were nonsynonymous.

## Data Analysis

All analyses, except for deep sequencing analysis, were performed using GraphPad Prism. For survival analysis, Kaplan-Meier survival curves were analyzed by the log-rank test. Unpaired Student’s t-test was used to determine significant differences in virus titers, crown-rump length, and viral loads. Fisher’s exact test was used to determine differences in rates of normal vs. abnormal concepti.

## Data Availability

Primary data that support the findings of this study are available at the Zika Open-Research Portal (https://zika.labkey.com). Zika virus sequence data have been deposited in the Sequence Read Archive (SRA) with accession code SRP150883. The authors declare that all other data supporting the findings of this study are available within the article and its supplementary information files, or from the corresponding author upon request.

## Acknowledgements

We thank Jody Peter, Victoria Leskinen, and Michelle Muhasky for maintenance of the *Ifnar1-/-* colony and assistance with timed-matings. Useful discussion with David O’Connor is greatly appreciated. Funding for this project came from National Institutes of Health grants R21AI131454, R01AI132563, R56AI132563, and start-up funds from the University of Minnesota Department of Veterinary and Biomedical Sciences to MTA; and National Institutes of Health grants R01AI067380 and R21AI125996 to GDE. RAM is supported in part by NSF training grant DGE-1450032 and by a Kirschstein National Research Service Award Individual Fellowship F31AI134108. The publication’s contents are solely the responsibility of the authors and do not necessarily represent the official views of the NCRR or NIH.

## Author Contributions

M.T.A., T.C.F., A.S.J., and G.D.E. designed experiments. A.S.J., T.C.F., and M.T.A. analyzed data and drafted the manuscript. G.D.E. and R.A.M. constructed the Zika virus infectious clone and engineered the mutant virus. R.A.M. and G.D.E. performed the *in vitro* viral replication experiments. A.S.J., M.T.A., and L.R.G. performed the mouse experiments. R.V.M., M.R.S., C.M.C., and S.L.O. developed and performed the deep sequencing pipeline. A.S.J. and L.R.G. performed plaque assays. A.S.J., A.M.W., S.R., M.R.S. and C.M.C. performed viral load assays. H.A.S., A.M., M.F., and C.H. performed histological sectioning and analyses. M.T.A., J.E.O., S.L.O, G.D.E., and T.C.F contributed space and reagents.

## Author Information

Reprints and permissions information is available at www.nature.com/reprints.

The authors declare no competing financial interests.

Correspondence and requests for materials should be addressed to

